# Implications of simplified linkage equilibrium SNP simulation

**DOI:** 10.1101/025619

**Authors:** S. Hong Lee

## Abstract

In a recent paper published in PNAS (Golan et al. 2014)^1^, residual maximum likelihood (REML) seriously underestimated genetic variance explained by genome-wide single nucleotide polymorphism when using a case-control design. It was concluded that Haseman–Elston regression (denoted as PCGC in their paper) should be used instead of REML. Their conclusions were based on results from simplified linkage equilibrium SNP simulation (SLES), which the authors acknowledged may be unrealistic. We found that their simulation, SLES, unrealistically inflated the correlation between the eigenvectors of the genomic relationship matrix and disease status to values that are rarely observed in real data analyses. With a more realistic simulation that the authors failed to carry out (as they noted in their paper), we showed that there was no such inflated correlation between the eigenvectors of the genomic relationship matrix and disease status. Because REML uses the eigensystem of covariance structure, the inflated correlation artefactually constrained its estimates. We compared SNP-heritabilities from SLES and a more realistic simulation, showing that there was a substantial difference between the REML estimates from the two simulation strategies. Finally, we presented that there was no difference between REML and PCGC in real data analyses. This pattern from real data results differed strikingly from the pattern in the simulation study of Golan et al.^1^. One needs to be cautious of results drawn from SLES.

---

Recently, Golan and Rosset (2013)^2^ and Golan et al. (2014)^1^ reported that REML seriously underestimates SNP-heritability when using a case-control design. Their conclusions were based on results from simplified linkage equilibrium SNP simulation (SLES), which the authors acknowledged may be unrealistic.

We simulated case-control data using the liability threshold model^1; 3^, based on a real GWAS of 800K SNPs from 64,000 samples, i.e. a genome-wide linkage disequilibrium SNP simulation (GLDS). Our simulation used a population disease risk of K = 0.01 and proportion of cases in the sample of P = 0.5 (therefore, there were 640 cases and 640 controls in the estimation analyses). A random 10,000, 1,000, or 100 SNPs across the genome were selected as risk loci. The genomic relationship matrix (GRM) was based on all the SNPs. For comparison, the SLES (without real GWAS data) was used, following Golan et al. (2014)^1^ where the GRM was calculated only from the risk SNPs that are independent from each other (see Supplementary Note).

In Figure 1, we show that SLES unrealistically inflates the correlation between the eigenvectors of the GRM and disease status compared to GLDS (Figure 2) or that inferred from real data (e.g., Figure S1 in Gusev et al. 2014^4^). The artefactual correlation between the eigenvectors and disease status caused the inaccuracy of the REML estimates. The bias depends on the ratio of the number of indiviudals (N) to the number of risk SNPs (M; Figure 1). Unlike REML, a sophisticated approach, Haseman–Elston regression (referred to as PCGC by Golan et al.^1^) does not use the eigensystem of covariance structure; therefore SLES does not affect the PCGC estimate^1^. With GLDS, the REML estimates were stable and close to the true value regardless of the value of N/M (Figure 3). With SLES, the REML estimates were severely biased with increasing value of N/M (Figure 3).

**Figure 1.**
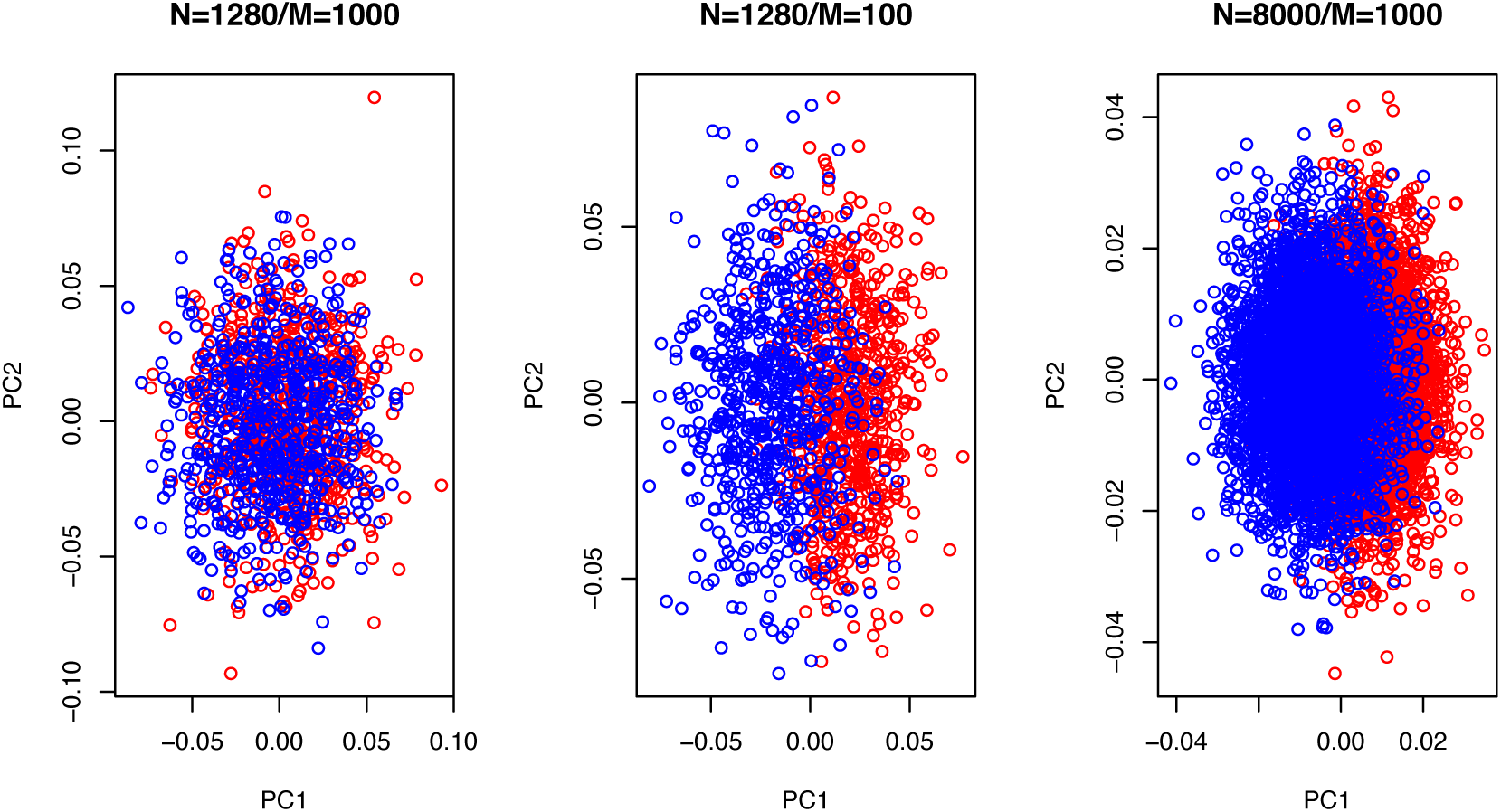
With the SLES simulation, the association between the eigenvectors and case-control status is unrealistically inflated when the value for N (# indiviudals) / M (# SNPs) increases. The correlation between the first principal component and disease status is 0.14, 0.70 and 0.63 with the value for N/M=1.3, 13 and 8, respectively. Population disease risk of K=0.01 and proportion of cases in the sample of P=0.5 were used. Red=cases and blue=controls.

**Figure 2.**
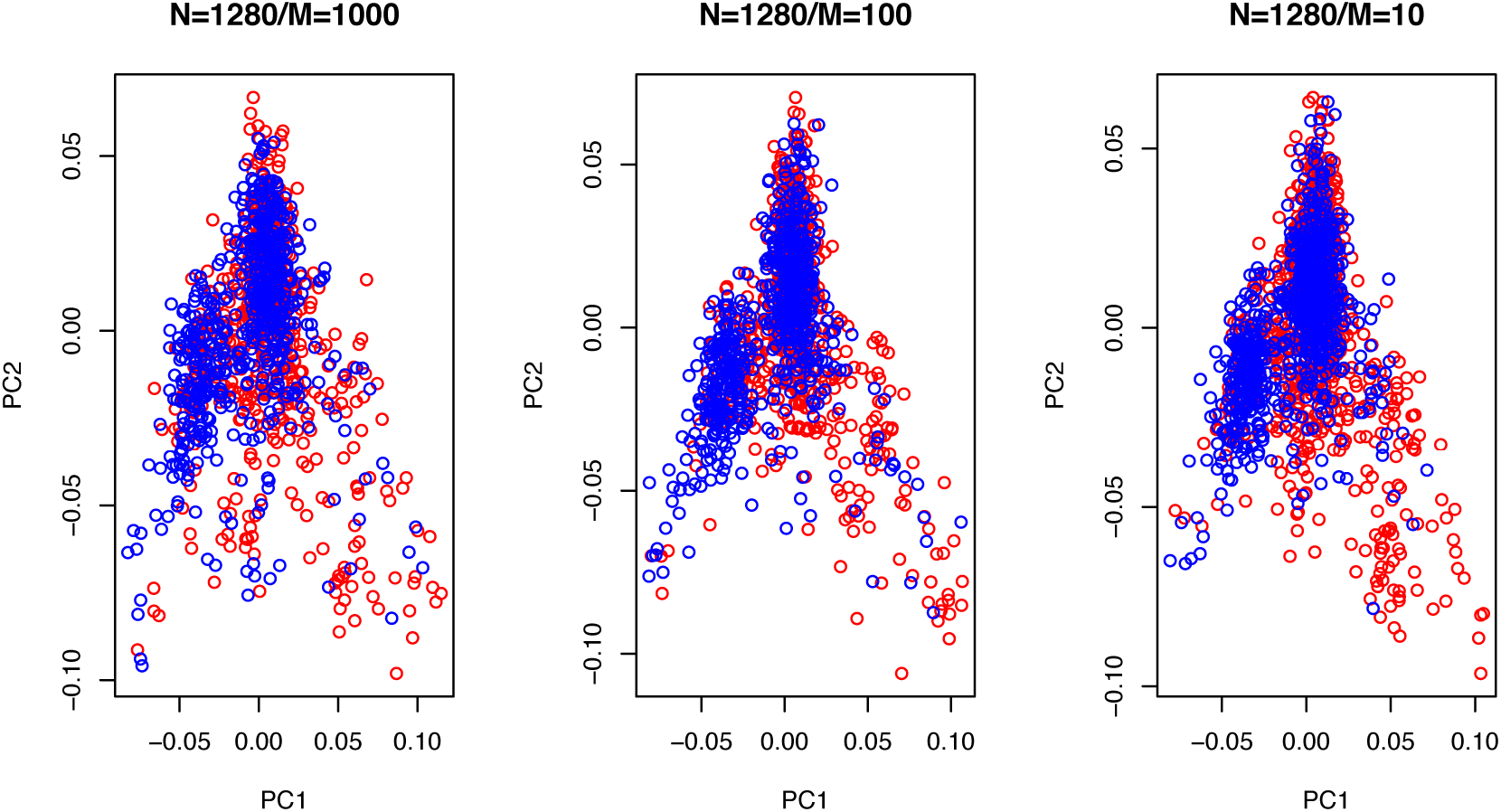
With the GLDS simulation, the association between the eigenvectors and case-control status is negligible, reagrdless of the values for N/M, i.e. more realistic compared to the SLES. The correlation between the first principal component and disease status is 0.04, 0.03 and 0.06 with the value for N/M=1.3, 13 and 130, respectively (GLDS could not simulate N=8000, since a GWAS data set of ∼400,000 individuals would be needed, instead we tested it with an extreme with M=10, i.e. N/M=130). Population disease risk of K=0.01 and proportion of cases in the sample of P=0.5 were used. Red=cases and blue=controls.

**Figure 3.**
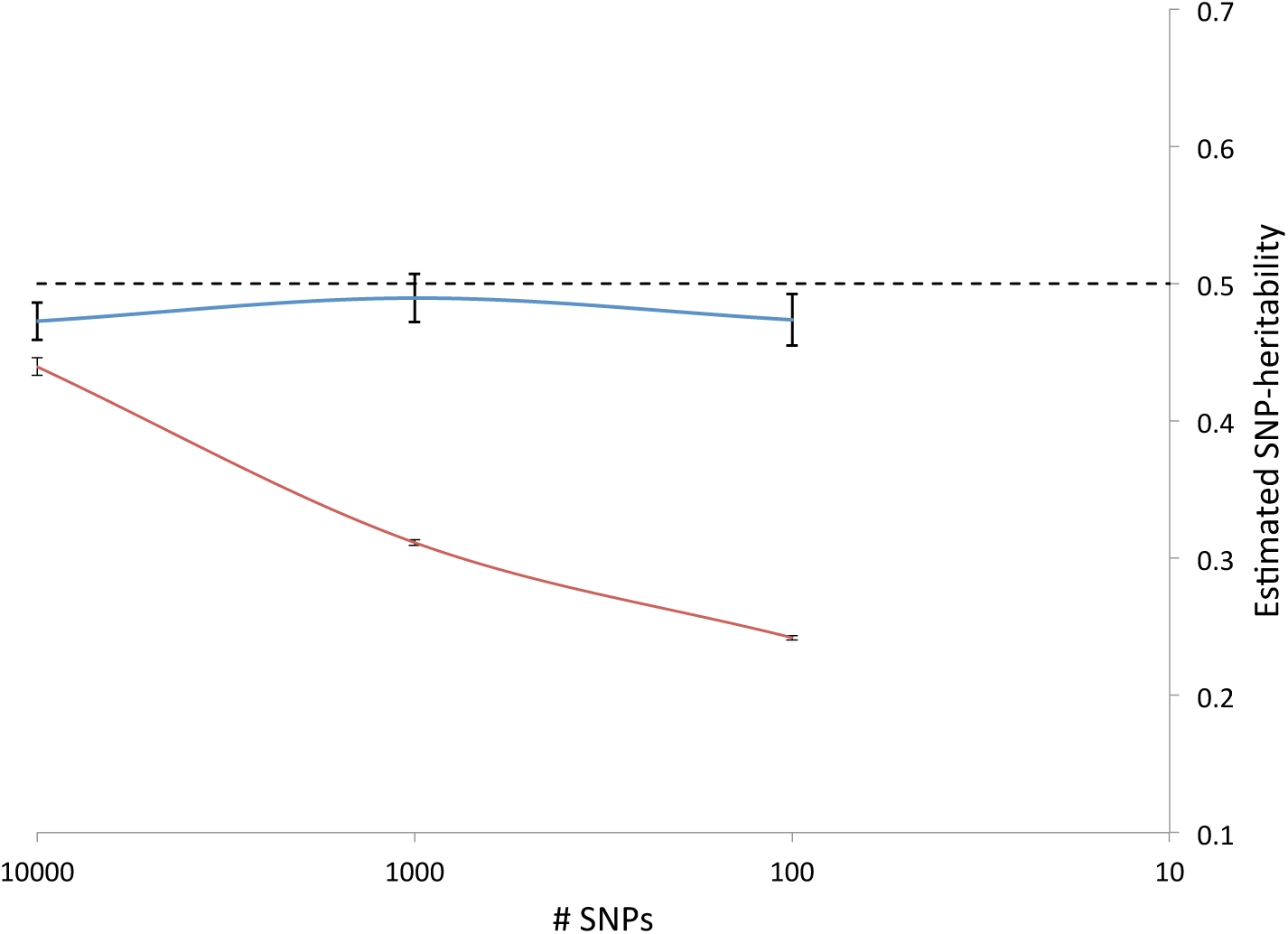
The average of estimated SNP-heritability and empirical SE bar of the mean estimate from REML with GLDS simulation (blue line) and SLES simulation (red line) over 50 replicates. The true simulated SNP-heritability is 0.5.

We considered results from real data analyses^4^ and plotted published SNP-heritability estimates against the sample size for nine diseases (Figure 4). There was no difference between REML and PCGC, regardless of sample size, which was strikingly different from Figure 2B in Golan et al.^1^ We also show estimation errors for the nine diseases assuming that the PCGC estimates are the true values (Figure 5), which were again dramatically different from results in Figure S4 from Golan et al. (2014)^1^.

**Figure 4.**
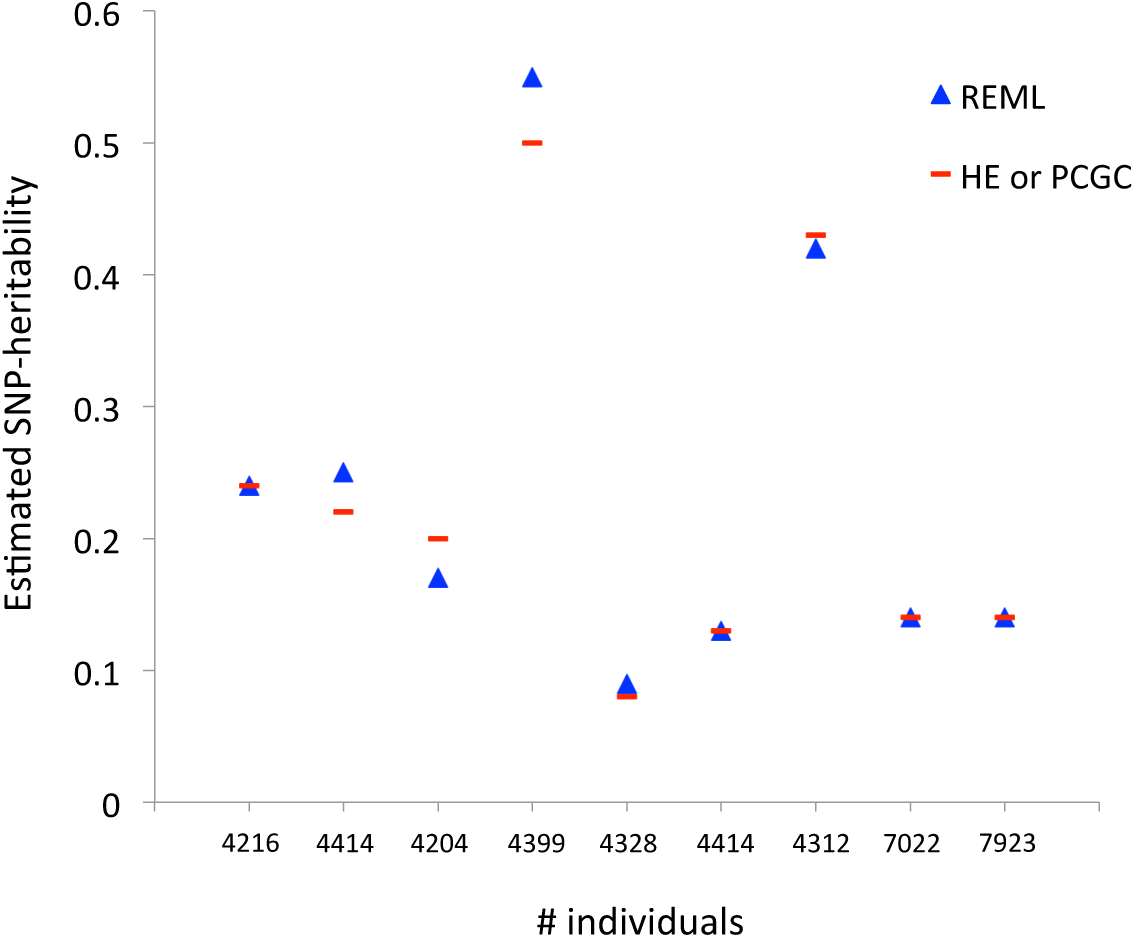
Estimated SNP-heritability from REML and PCGC with real data analyses (to be compared to Figure 2B in Golan et al. 2014). We excluded two diseases that had highly confounded population structure (see Figure S1 in Gusev et al. 2014^4^). HE: Haseman-Elston regression.

**Figure 5.**
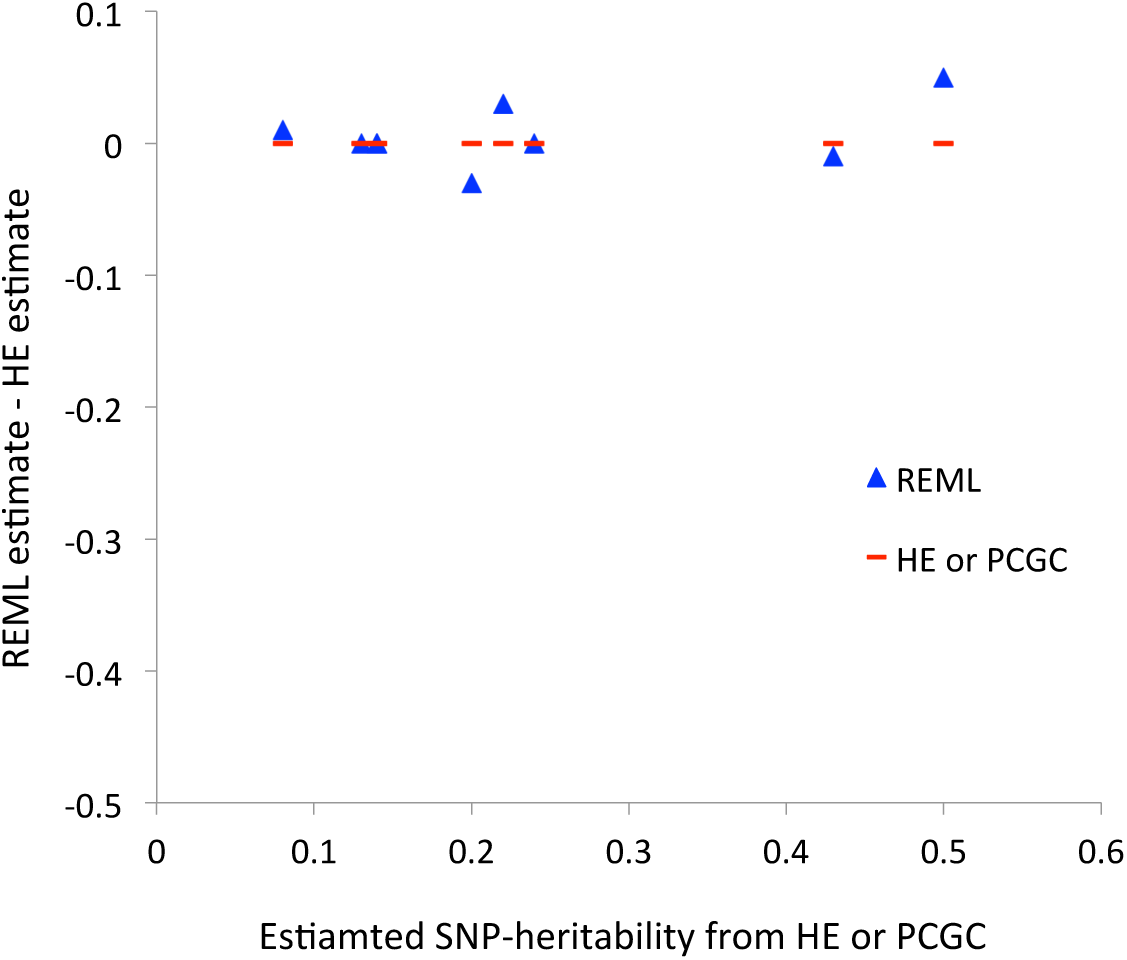
Estimation error assuming that PCGC estimate is true value (to be comapred to Figure S10 in Golan et al. 2014).

There are a large number of unknown confounding factors, such as artefact batch effects, in real case-control data. A number of studies have emphasised that such unknown factors can inflate estimates and should be controlled by stringent QC^5-7^. In addition to stringent QC (MAF, missingness, H-W test, missingness difference, relatedness cut-offs), Gusev et al. (2014)^4^ additionally performed five rounds of outlier removal whereby all individuals more than 6 SDs away from the mean along any of the top 20 eigenvectors were removed and all eigenvectors were recomputed. Moreover, they fit 20 principal components (PCs) into their SNP-heritability estimation to correct for population admixture. Less stringent QC and/or less number of PCs fitted in the model might result in less effective control of population admixture as well as artifact batch effects.

In derivation of the correction factor for case-control ascertainment bias, Lee et al. (2011)^3^ used a simulation from a multivariate normal distribution based on a predefined relationship matrix. In real data analyses, the true relationship matrix is not known but can be approximated from genotypes, i.e. GRM pairwise estimator is unbiased under linkage disequilibrium; that is, the expectation of the estimator for each SNP is the kinship in the identical-by-descent (IBD) fraction sense^8^, and therefore so is the estimate averaged over multiple SNP’s. SLES ignores the concept of linkage disequilibrium, IBD and coalescence. We urge researchers to use a more realistic genetic model (e.g., GLDS at least) in their simulation strategies and to be cautious of results drawn from SLES^1; 9^.

